# Transcriptomic profiling of human effector and regulatory T cell subsets identifies predictive population signatures

**DOI:** 10.1101/2020.05.13.093567

**Authors:** Barbara Höllbacher, Thomas Duhen, Samantha Motley, Maria M. Klicznik, Iris K. Gratz, Daniel J. Campbell

## Abstract

After activation, CD4^+^ T helper (Th) cells differentiate into functionally specialized populations that coordinate distinct immune responses and protect against different types of pathogens. In humans, these effector and memory Th cell subsets can be readily identified in peripheral blood based on their differential expression of chemokine receptors that govern their homeostatic and inflammatory trafficking. Foxp3^+^ regulatory T (Treg) cells can also be divided into subsets that phenotypically mirror each of these effector populations, and share expression of key transcription factors and effector cytokines. In this study, we performed comprehensive transcriptional profiling of 11 phenotypically distinct Th and Treg cell subsets sorted from peripheral blood of healthy individuals. Despite their shared phenotypes, we found that mirror Th and Treg subsets were transcriptionally dissimilar, and that Treg cell populations showed limited transcriptional diversity compared to Th cells. We identified core transcriptional signatures shared across all Th and Treg cell populations, and unique signatures that define each of the Th or Treg populations. Finally, we applied these signatures to bulk Th and Treg RNA-seq data and found enrichment of specific Th and Treg cell populations in different human tissues. These results further define the molecular basis for the functional specialization and differentiation of Th and Treg cell populations, and provide a new resource for examining Th and Treg specialization in RNA-seq data.

## Introduction

The outcome of adaptive immune responses is dictated in large part by the activity of CD4^+^ T helper (Th) cells. Th cells differentiate into functionally specialized populations that through secretion of effector cytokines coordinate the activities of immune and stromal cells to control different types of pathogens and dangerous toxins (Sallusto, 2016). Thus IFN-γ-producing Th1 cells are essential for control of intracellular pathogens, Th2 cells that produce IL-4, IL-5 and IL-13 mediate protection against helminth infection and poisonous venoms, and Th17 cells that produce IL-17 and IL-22 provide immunity to extracellular bacteria and fungal pathogens such as *Staphylococcus aureus* and *Candida albicans*. More recently described Th populations include Th22 cells (Duhen et al., 2009; Trifari et al., 2009), which produce IL-22 but not IL-17, and appear to function specifically within the skin during tissue-repair responses, and Th1/17 cells that share key features with both Th1 and Th17 cells including co-production IFN-γ and IL-17, and are enriched in cells producing GM-CSF (Annunziato et al., 2007; Duhen and Campbell, 2014). In human blood, these Th subsets can be readily identified based on their differential expression of chemoattractant receptors and adhesion molecules that control their specific migration to distinct inflammatory sites and likely sites of pathogen entry.

Although essential for protection against infection, dysregulated Th cell responses are pathogenic in immune-mediated diseases. These include organ-specific autoimmune diseases such Type-1 diabetes (Th1), psoriasis (Th17, Th22), and multiple sclerosis (Th1/17), as well as allergic hypersensitivities and asthma (Th2). The activities of Th cells are restrained by T regulatory (Treg) cells, a subset of CD4^+^ T cells that constitutively expresses the IL-2 receptor alpha chain CD25 and the transcription factor Foxp3. Treg cells dampen the activation and function of Th cells via multiple mechanisms, including production of anti-inflammatory cytokines such as IL-10, IL-35 and TGF-β, CTLA4-mediated blockade of T cell co-stimulation, sequestration of effector cytokines, metabolic disruption of Th cells, competition for peptide:MHC complexes, and direct cell lysis (Shevach, 2018). Importantly, we and others have shown that Treg cells are not homogenous, but instead like Th cells can be divided into diverse subsets (Campbell and Koch, 2011). These phenotypically ‘mirror’ each of the major Th populations, share expression of lineage-defining transcription factors such as T-bet and RORγt, and can even produce effector cytokines such as IFN-γ and IL-17 (Dominguez-Villar et al., 2011; Duhen et al., 2012). These shared features suggest that Th and Treg cells may be more similar than previously appreciated. We also showed that production of the immunosuppressive cytokine IL-10 was limited to Th1- and Th17-like Treg cells, suggesting that different Treg cell subsets may employ specialized immunoregulatory mechanisms to modulate different types of inflammatory responses.

To address these issues, we performed comprehensive transcriptional profiling of 11 distinct CD4^+^ T cell subsets sorted from the peripheral blood of 3 healthy donors. We found that although they shared many phenotypic features, mirror Th and Treg cell populations were transcriptionally divergent, and that the Treg cell populations showed limited transcriptional diversity compared to Th cells. We further identified core transcriptional signatures shared across all Th or Treg cells, as well as unique signatures that define each of the Th or Treg cell populations. Finally, we applied these signatures to RNA-seq data from bulk Th and Treg cells to show enrichment of specific Th and Treg cell populations in healthy and diseased human tissues.

## Materials and Methods

### Flow cytometric sorting

Blood samples were obtained from healthy donors participating in the Benaroya Research Institute Immune-Mediated Disease Registry. Informed consent was obtained from all subjects according to institutional review board-approved protocols at Benaroya Research Institute and following the Declaration of Helsinki. CD4^+^CD25^high^ Treg cells were enriched from PBMCs after staining with PE-cyanine 5 (PE-Cy5)-labeled anti-CD25 Ab (BioLegend), followed by positive selection using anti-PE and anti-Cy5 microbeads (Miltenyi Biotec). On the negative fraction, CD4^+^CD25^−^ Th cells were purified by positive selection with CD4-specific microbeads (Miltenyi Biotec). Memory T-cell subsets were sorted to more than 97% purity as CD4^+^CD45RA^−^ RO^+^CD127^+^CD25^−^ using APC-780-conjugated anti-CD45RA Ab (eBioscience), Alexa 700-conjugated anti-CD45RO Ab (BioLegend), v450-conjugated anti-CD127 Ab (BD Bioscience), PE-Cy5-conjugated anti-CD25 Ab (BioLegend) and Qdot655-conjugated anti-CD4 Ab (eBioscience). Abs used for sorting of memory Th and Treg cell subsets were: PE-Cy7-conjugated anti-CCR6 Ab (BioLegend), PE-conjugated anti-CCR10 Ab (R&D Systems), PerCP/Cy5.5-conjugated anti-CCR4 Ab (BioLegend), and Alexa Fluor 488-conjugated anti-CXCR3 Ab (BD Bioscience). Cells were sorted with a FACSAria II (BD Biosciences).

### Library construction and RNA-seq

RNA-seq libraries were constructed from up to100 ng of total RNA using the TruSeq RNA Sample Prep Kit v2 (Illumina). Libraries were clustered on a flowcell using the TruSeq Paired-end Cluster Kit, v3 using a cBot clustering instrument (Illumina), followed by paired-end sequencing on a HiScanSQ (Illumina) for 50 cycles in either direction. After the run was completed, the reads were demultiplexed and FASTQs were generated for each sample using CASAVA.

### RNA-seq Analysis

Base-calling was performed automatically by Illumina real time analysis software and demultiplexing was performed with the program Casava. One 3’-end base was removed from all reads, followed by quality-based trimming from both ends until minimum base quality for each read was >= 30. Tophat aligned reads to GRCh38, using Ensembl annotation release number 77 and the read counts per Ensembl gene ID were computed with featureCounts. Sequencing, alignment, and quantitation metrics were obtained for FASTQ, BAM/SAM, and count files using FastQC, Picard, TopHat, Samtools, and htseq-count. All samples passed QC with mapped reads with duplicates > 80%, median CV coverage < 0.8 and total fastq reads of > 5 Mio. Protein coding transcripts with a minimum of 1 CPM in at least 5% of the total number of libraries were retained and the ensembl gene IDs mapped to HGNC gene symbols. Count data was normalized using edgeR’s TMM (Robinson and Oshlack, 2010). For linear modelling we computed a coefficient for each subset with the Naïve population as reference and accounted for donor variation as random factor using the R package limma (Ritchie et al., 2015). To determine differences between the Th and Treg cells that are common for all subsets, we set up pairwise contrasts between mirror Th and Treg subsets and examined the overlap. For visualizations such as PCA and expression plots, this is approximated with the limma function removebatcheffect. Cluster means were computed with the kmeans function for 3 total clusters, transformed into PCA space and added to the PCA plot. For gene expression dotplots the base of 2 was raised to the power of rembatch corrected values (which were originally taken from output of the voomwithqualityweights function) in order to unlog them. P values for the venn diagrams of DE genes between Tregs and their respective Th counterpart were adjusted using global BH adjustment of FDR=0.05 and summarized with the limma::decideTest function including a cutoff of absolute log2 fold change > 1. Heatmaps are based on log-transformed expression values, are z-scaled by rows and were plotted using the R package ComplexHeatmap (Gu et al., 2016). Euclidean distances between all samples were computed with default settings from the stats::dist() function and plotted as a heatmap with manually defined sample order. To compare the heterogeneity of Th and Treg populations, we recomputed the distances of Treg to Treg samples, Th to Th samples and Treg to Th samples separately. The results are summarized and plotted as density estimates using ggplot. Unique subset signatures were determined by making contrasts of a given subset with all other Th or Treg populations (excluding Th1/17 cells), and genes that were differentially expressed at adj. p value < 0.05 between a given subset and all other subsets were termed to be part of a population signature. To evaluate the relative expression of signature genesets in other samples, we rank ordered all genes for a given samples and computed the mean rank for the significantly upregulated genes in each signature. Gene expression data of IFN-γ- and IL10-producing Treg populations from the public dataset GSE116283 were accessed through GEO and filtered for genes with CPM >1 in at least 25% of all samples that were also present in our dataset. Gene expression matrix and series matrix of breast cancer and PBMC samples were retrieved through the GEO accession number GSE89225. The samples run on the Iontorrent platform were downloaded and assigned to their corresponding HGNC gene keys. Gene expression data was filtered for protein coding genes that were also expressed in our dataset and those with CPM > 1 in at least 25% of the samples.

### Data accessibility

RNA-seq data generated for this study are available from GEO as SuperSeries GSE149090.

## Results

### RNA-seq analysis of human CD4^+^ T cell populations

To comprehensively profile human CD4^+^ Th and Treg cells, we performed RNA-seq on 11 distinct CD4^+^ T cell populations sorted directly from peripheral blood of 3 healthy individuals on the basis of surface markers and chemokine receptors as previously described (Duhen et al., 2012) (Fig 1). Th cells were identified as CD127^+^CD25^−^, and were sorted into naïve (CD45RA^+^), as well as Th1 (CD45RA^−^CXCR3^+^CCR6^−^), Th17 (CD45RA^−^CXCR3^−^CCR6^+^CCR4^+^CCR10^−^), Th1/17 (CD45RA^−^CXCR3^+^CCR6^+^), Th2 (CD45RA^−^CXCR3^−^CCR6^−^CCR4^+^) and Th22 (CD45RA^−^CXCR3^−^CCR6^+^CCR4^+^CCR10^+^) cell fractions. Treg1, Treg17, Treg1/17, Treg2 and Treg22 populations were sorted within the CD127^−^CD25^+^ Treg cells using the same markers (Fig 1). Expression of genes encoding the surface receptors used for cell sorting faithfully segregated into the various Th and Treg transcriptional profiles, and analysis of genes encoding key transcription factors associated with Treg cells (*FOXP3*), or with Th1/Treg1 (*TBX21*), Th17/Treg17 (*RORC*) and Th2/Treg2 (*GATA3*) subsets were expressed as expected within the appropriate cell populations (Fig 2A). Thus, the sorting strategy we employed properly isolated functionally specialized Th and Treg cell populations, and our data provide a novel and comprehensive transcriptional analysis of *in vivo* differentiated Th and Treg cells.

**Figure 1:**
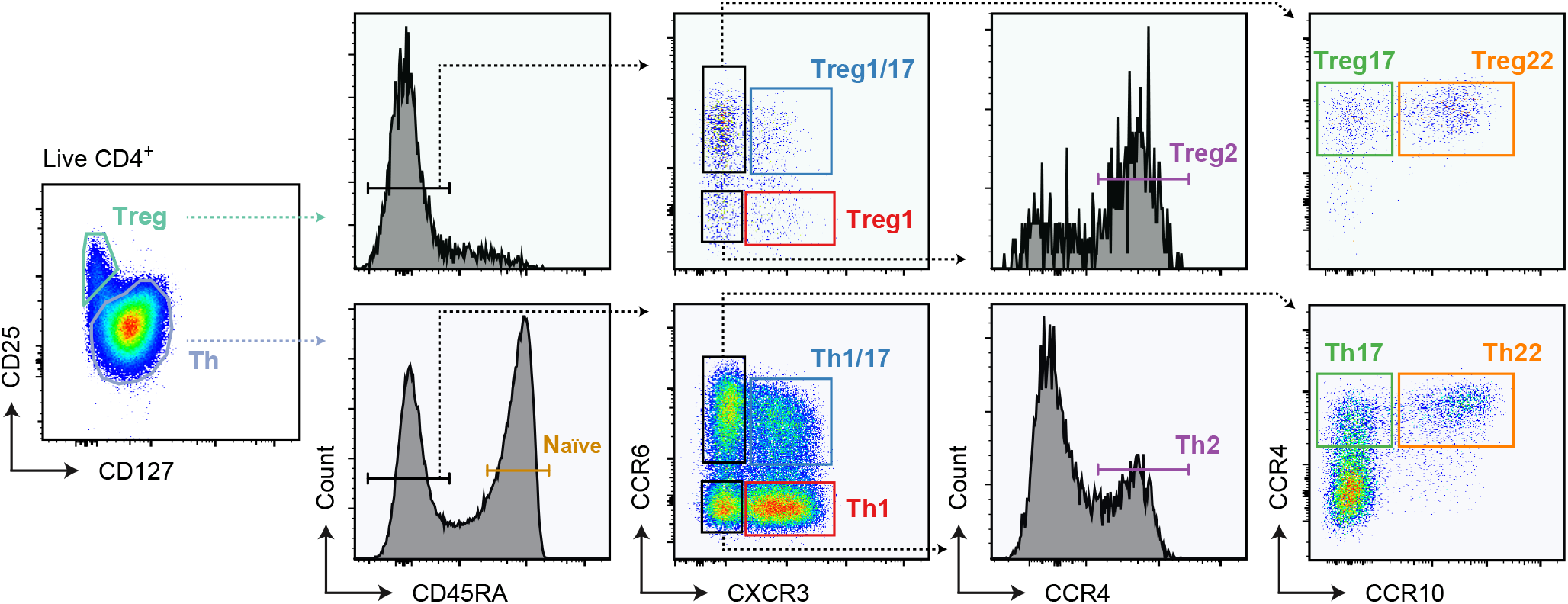
Gating strategy used to sort Th, Treg and Naive cell populations. Representative flow cytometric analysis of gated CD4^+^ peripheral blood mononuclear cells showing gating strategy used to sort each of the 11 T cell populations for RNA-seq analysis.

**Figure 2:**
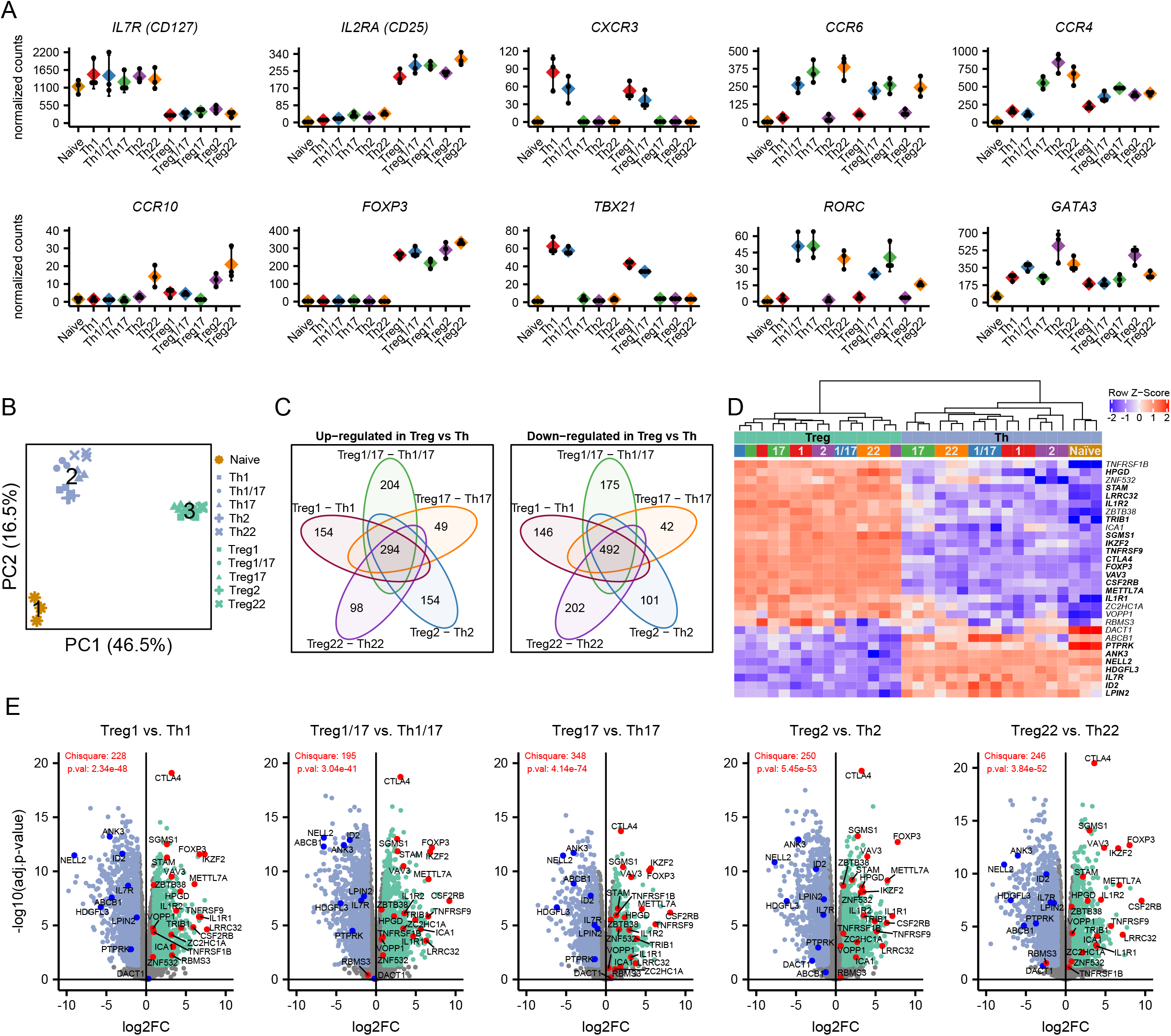
Identification of core Th and Treg cell signatures. **A)** Expression analysis of the indicated chemokine receptors and transcription factors that define the different T cell populations. Values represent batch-effect corrected counts per million. **B)** Principal component analysis of all samples run on all expressed genes. Cluster centers representing naïve T cells (1), Th cells (2) and Treg cells (3) returned by k-means clustering. **C)** Venn diagrams showing the overlap in genes that were significantly (adjusted p value < 0.05 and absolute log2 fold change > 1) up-regulated (left) or down-regulated (right) when comparing Tregs with their mirror Th counterpart as indicated. Numbers indicate genes that were up-/down-regulated in all comparisons (core of the venn diagrams), or genes that were differentially expressed in only one comparison. **D)** Heatmap showing expression of 31 genes identified by Pesenacker et al. as activation independent Treg markers in the indicated samples. Bolded genes were significantly differentially expressed in all comparisons of mirror Th and Treg populations. **E)** Volcano plots showing pairwise comparisons of mirror Th and Treg cell populations. Genes identified by Pesenacker et al. as higher in Treg cells are highlighted in red, those higher in Th cells in blue. Chisquare test statistic and p-values indicate whether the sets of up- and down-regulated identified by Pesenacker et al. are differentially distributed in the indicated Treg vs. Th comparison.

### Th and Treg cells are transcriptionally distinct

Using principle component analysis (PCA), we compared the overall transcriptional profile of all Th and Treg subsets (Fig 2B). Kmeans clustering was used to identify three distinct clusters, which corresponded to samples from naïve Th cells (cluster 1), memory Th cells (cluster 2) and Treg cells (cluster 3). Thus, despite their phenotypical similarity and shared expression of key lineage-defining transcription factors, mirror Th and Treg cell populations (e.g., Th1 and Treg1 cells) are transcriptionally distinct and more closely resemble other Th and Treg cells. To define core Th and Treg transcriptional signatures, we performed pairwise comparisons between each of the individual Th and Treg cell mirror pairs, and identified 294 genes that were significantly higher (log_2_ fold change>1, adjusted p-value<0.05) in all Treg cell populations, and 492 that were more highly expressed in all Th populations (Fig 2C). Analysis of these gene sets revealed many genes previously identified as differentially expressed in Treg vs. Th (Bhairavabhotla et al., 2016; Pesenacker et al., 2016) (Supplementary Table 1). These included *FOXP3*, *CTLA4*, *ENTPD1* (CD39), *IKZF2* (Helios), and *TNFRSF9* that are preferentially expressed in Treg cells, and *BHLHE40, CD40LG, ID2*, and *IL2* that are more highly expressed in Th cells.

Because Treg cells are largely specific for either auto-antigens or for components of the microbiome present on barrier surfaces, they are subject to chronic stimulation and many of the genes previously associated with Treg cells are activation-induced genes that are not actually Treg cell specific (Pesenacker et al., 2016). Consistent with this, among genes more highly expressed in Treg cells, several were genes we previously found to be downregulated in subjects treated with the co-stimulation blocking drug abatacept (CTLA4-Ig) (Glatigny et al., 2019), including *ARHGAP11A*, *CENPE*, *DUSP4*, *ERI1*, *GXYLT1*, *HELLS*, *NUSAP1*, *PMAIP1*, *SGMS1*, and *TOP2A* (Supplementary Table 1). To account for these and other activation-induced genes, Pesenacker et al. compared the transcriptomes of resting and activated Th and Treg cells, and identified a 31-gene ‘activation-independent’ Treg cell signature (Pesenacker et al., 2016). Of these, 21 were differentially expressed in all 5 Th/Treg cell comparisons, and of the remaining genes 3 were significant in 4 out of 5 comparisons (Fig 2D,E).

In our PCA analysis (Fig 2B), all Treg cell populations clustered tightly together (cluster 3), whereas there was substantially more spreading within the cluster of memory Th cell populations (cluster 2), indicating that overall Treg cells are transcriptionally less diverse than the Th cells. To further examine this, we calculated the Euclidian distance of all samples relative to each other in order to quantitatively assess their overall similarity (Fig 3A, B). As evidenced in the PCA, Treg and Th subsets were highly divergent, with a mean Euclidian distance in these comparisons of 129.7. The Euclidian distance comparing all the Th cell samples to each other had a bi-modal distribution, with a peak of highly similar samples corresponding to comparisons between the same Th populations isolated from the 3 donors (e.g., comparing all Th1 samples, mean Euclidian distance 59.06), and a second significantly more distant peak corresponding to comparisons between the different types of Th cell populations (mean Euclidian distance 82.44, p<2×10^−16^, Welch two sample t-test). By contrast, we found that the Euclidian distance between different Treg populations was relatively uniform, and that the mean Euclidian distance between all of the Treg populations was significantly smaller than observed in the comparisons of the Th populations (mean Euclidian distance 69.71, p<2×10^−16^, Welch two sample t-test). Thus, despite their similar degree of phenotypic heterogeneity, the different Treg cell populations are transcriptionally less diverse than their Th counterparts.

**Figure 3:**
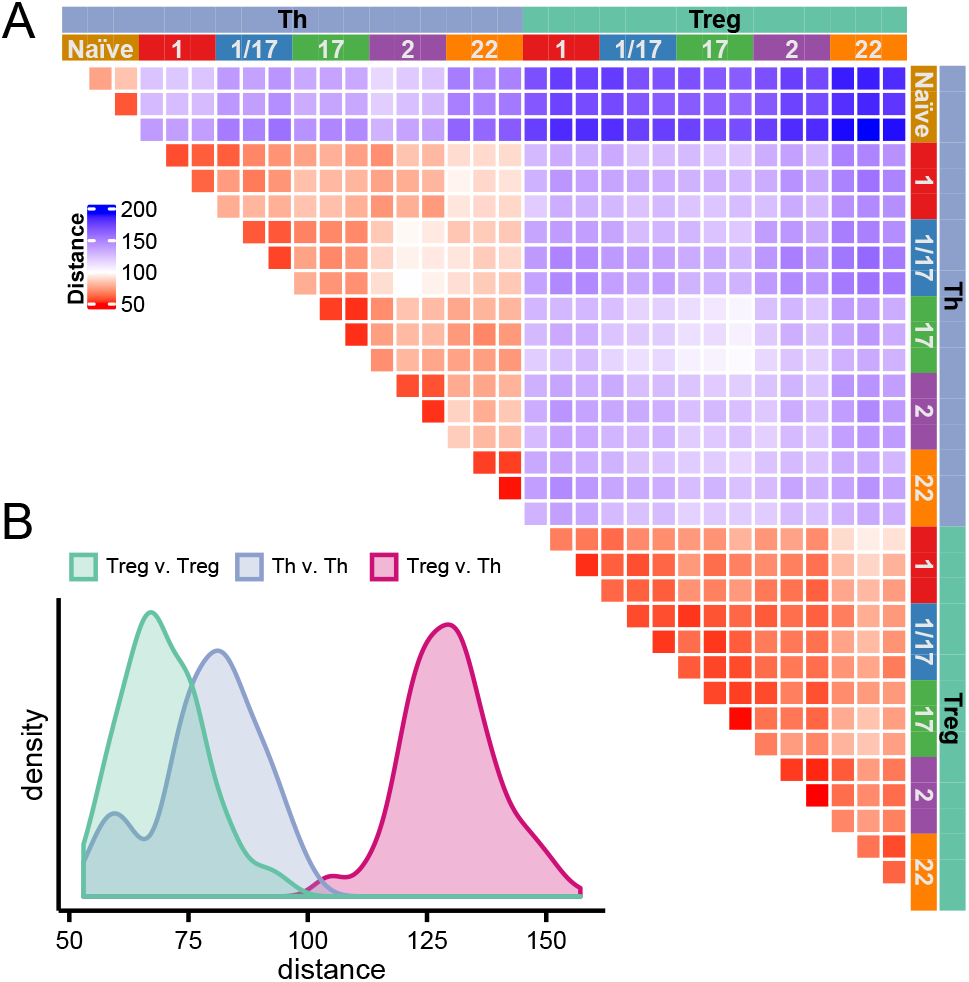
Treg cell subsets show limited transcriptional diversity. **A)** Heatmap representing the Euclidian distances (based on batch effect corrected expression values) in pairwise comparisons of all samples as indicated. **B)** Histograms showing the distributions of Euclidian distances in comparisons of all Treg vs Th, Th vs. Th, or Treg vs. Treg as indicated.

### Functional specialization of Treg cells

The differentiation of Th cells into specialized subsets is accompanied by acquisition of distinct effector functions that coordinate different types of immune responses. Similarly, a large number of effector mechanisms are used by Treg cells to dampen inflammation and prevent autoimmunity. These include production of anti-inflammatory cytokines, expression of co-inhibitory receptors, cytokine sequestration, modulation of pro- and anti-inflammatory metabolites, and direct target cell lysis. This raises the intriguing possibility that like Th cells, different Treg cell subsets may employ specialized immunoregulatory mechanisms to modulate different types of inflammatory responses. Indeed, we previously demonstrated that production of the anti-inflammatory cytokine IL-10 was restricted to Treg1 and Treg17 cells (Duhen et al., 2012). To further examine the functional specialization of Treg cells, we assessed expression of a set of key genes controlling Treg cell activity in different settings (Fig 4A). Consistent with our previous findings, *IL10* expression was largely limited to the Treg1, Treg17 and Treg1/17 populations, and these also selectively expressed the effector cytokines *IFNG* and *IL17A* which has previously been reported for human Treg cells. Treg1 cells also most highly expressed the co-inhibitory receptors *LAG3* and *HAVCR2* (TIM-3), and the cytolytic effectors *GZMA* and *GZMK*. By contrast, expression the TGF-β-activating molecule *LRRC32* (GARP) was highest in Treg 2 cells, the decoy IL-1 receptor *IL1R2* was most highly expressed by Treg 17 and Treg1/17 cells, and HPGD, which is used by Treg cells to degrade the pro-inflammatory prostaglandin PGE2 (Schmidleithner et al., 2019), was most highly expressed in Treg22 cells. Other genes implicated in Treg function including *TGFB1*, *ITGAV*, *ITGB8*, *IL12A* (a component of the anti-inflammatory cytokine IL-35), *CTLA4*, *PDCD1* (PD1), *TIGIT*, *PRF1*, and *ENTPD1* (CD39) did not show strong preferential expression in any Treg population, and *NT5E* (CD73) was barely detected in these *ex vivo* isolated Treg cells.

**Figure 4:**
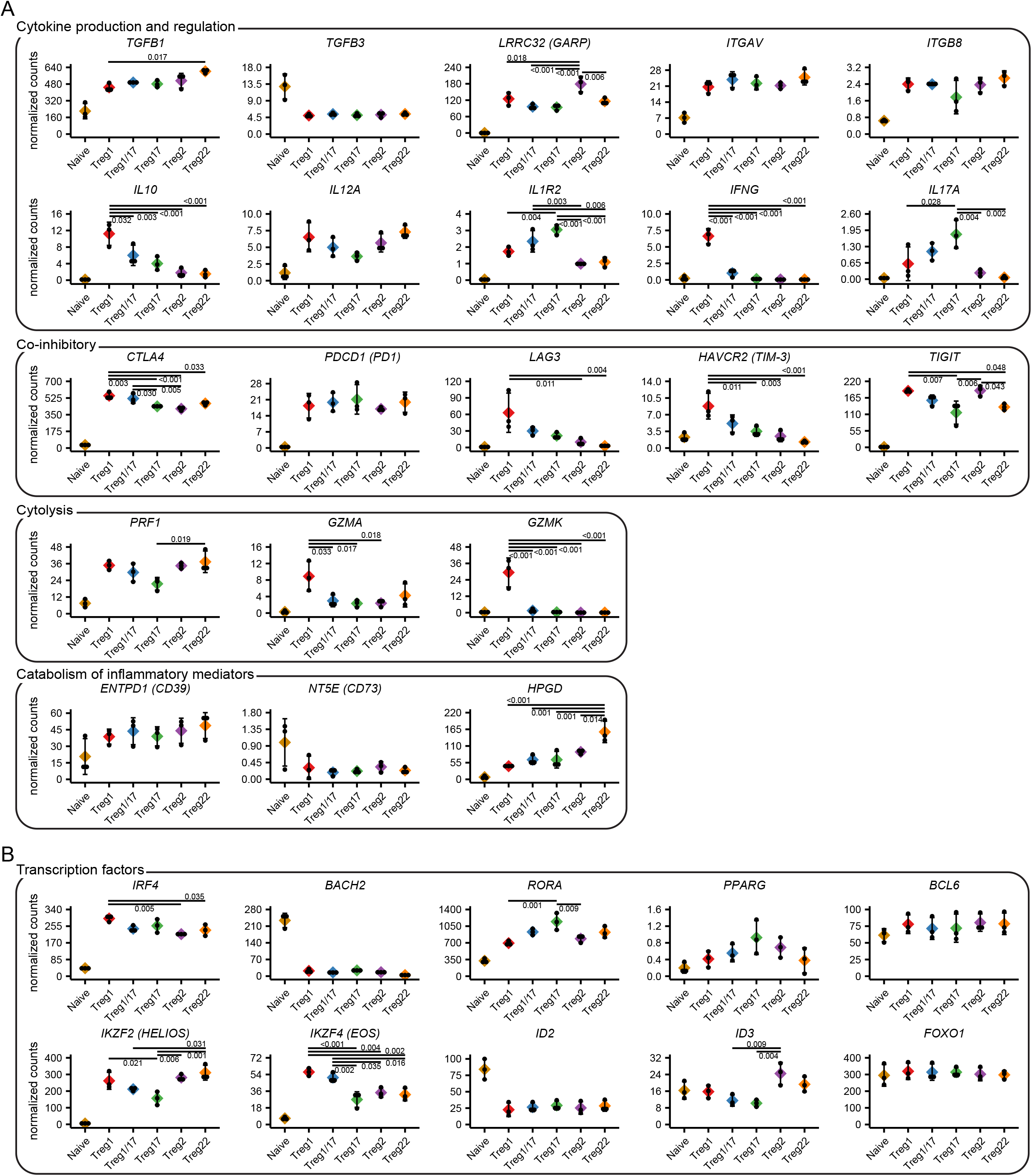
Functional and molecular specialization of Treg cell populations. **A)** Expression analysis of genes controlling different functions of Treg cells as indicated. **B)** Expression analysis of transcription factors implicated in Treg specialization. Values represent batch-effect corrected counts per million. Significance levels were determined by ANOVA followed by TukeyHSD for testing pairwise differences. Significant differences with the naïve samples are not highlighted.

Treg cell specialization depends on their selective expression of a set of transcription factors that alter their migration and function. In addition to T-bet (*TBX21*), GATA3 (*GATA3*) and RORγt (*RORC*) that promote Treg1, Treg2 and Treg17 differentiation, other transcription factors implicated in control of Treg cell function include *IRF4*, *BACH2*, *RORa*, *PPARG*, *BCL6*, *IKZF2* (Helios), *IKZF4* (Eos), *ID2*, *ID3*, and *FOXO1*. Indeed, *IRF4, RORA, IKZF2, IKZF4* and *ID3* were significantly differentially expressed between various Treg cell subsets, and this may contribute to their molecular specialization. Expression of *PPARG* was highest in Treg17 cells, but this did not reach statistical significance. By contrast, *BCL6* and *FOXO1* were expressed equally in all Treg populations whereas *ID2* and *BACH2* showed limited expression in all Treg populations relative to naïve T cells, and therefore these factors are unlikely to contribute to their phenotypic and functional heterogeneity.

### Identification of subset specific transcriptional signatures in Th and Treg cells

Although the functional specialization of Th cells was first described over 30 years ago, their complex phenotypes, specialized functions and pathways of differentiation are still being defined. Much of our understanding of human Th cell differentiation has been derived from cells differentiated *in vitro*, and these may differ substantially from cells differentiated *in vivo*. Therefore, to gain further insight into the functional specialization of Th cells, we identified the unique transcriptional signature for each Th subset based on genes significantly up- or down-regulated in comparison with all other Th subsets (Fig 5A). Because Th1/17 cells are a hybrid population with characteristics of both Th1 and Th17 cells (Annunziato et al., 2007), we excluded them from these comparisons in order to more readily define the Th1 and Th17 subset signatures. Analysis of these signatures revealed selective expression of genes that were previously implicated in Th cell specialization, localization and function including *IFNG, IL12RB2, TBX21, and EOMES* that are selectively expressed in Th1 cells; *IL12RB1* and *ABCB1* in Th17 cells; *GATA3*, *PTGDR2*, and *IL4R* in Th2 cells; and *ITGAE*, *CD9*, and *CD101* in Th22 cells. We also identified genes in each of the subset signatures that may differentially influence cell metabolism (decreased *HK2* expression in Th1 cells), cell signaling (increased *SOCS2* expression in Th17 cells), and cell function (increased *TNFSF11* (RANKL) expression in Th2 cells). Similarly, by comparing the transcriptional profiles of the Treg subsets (excluding Treg1/17 cells), we identified population-specific transcriptional signatures in Treg cells (Fig 5B). A full list of upregulated and downregulated genes in all Th and Treg population signatures is found in Supplementary Table 2.

**Figure 5:**
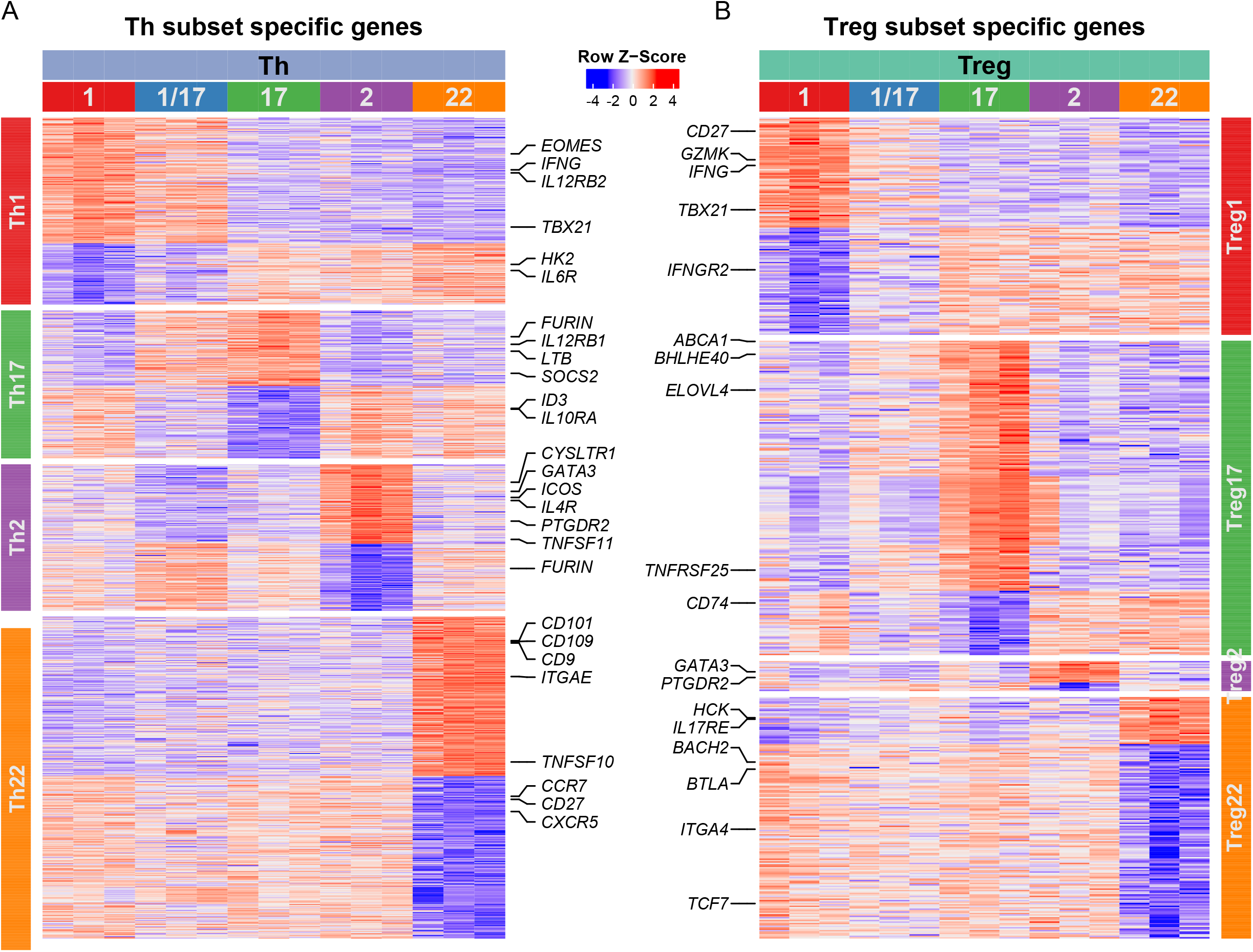
Identification of subset-specific transcriptional signatures in Th and Treg cells. Heatmaps representing expression of genes in each of the indicated cell samples (top labels) that were up-regulated or down-regulated (adjusted p value < 0.05 and absolute log2 fold change > 1) in each of the **A)** Th cell or **B)** Treg cell populations (side labels) as indicated. Select genes in each of the population signatures are highlighted.

To assess expression of the gene signatures between different populations, we ranked all genes in order of expression for each sample, and determined the mean genelist rank for each set of signature genes in all populations examined. For this, we focused on genes that were upregulated in each of the signatures and therefore positively identified a given population. This analysis demonstrated that the signatures of both the Th1 and Th17 populations were also shared in the Th1/17 cells (Fig 6A), consistent with the Th17/17 population having hybrid characteristics of both Th1 and Th17 cells. Analysis of the gene signatures of mirror Th and Treg populations revealed little overlap in the identity of the differentially expressed genes (Fig 6B). However, examining the mean genelist rank showed that in all cases, Th cell subset signatures were significantly enriched in the corresponding Treg cell subset, and that Treg subset signatures were significantly enriched in the corresponding Th populations (Fig 6C). Thus, despite their overall transcriptional dissimilarity, the Th and Treg cell populations do share some common phenotype-associated transcriptional programs.

**Figure 6:**
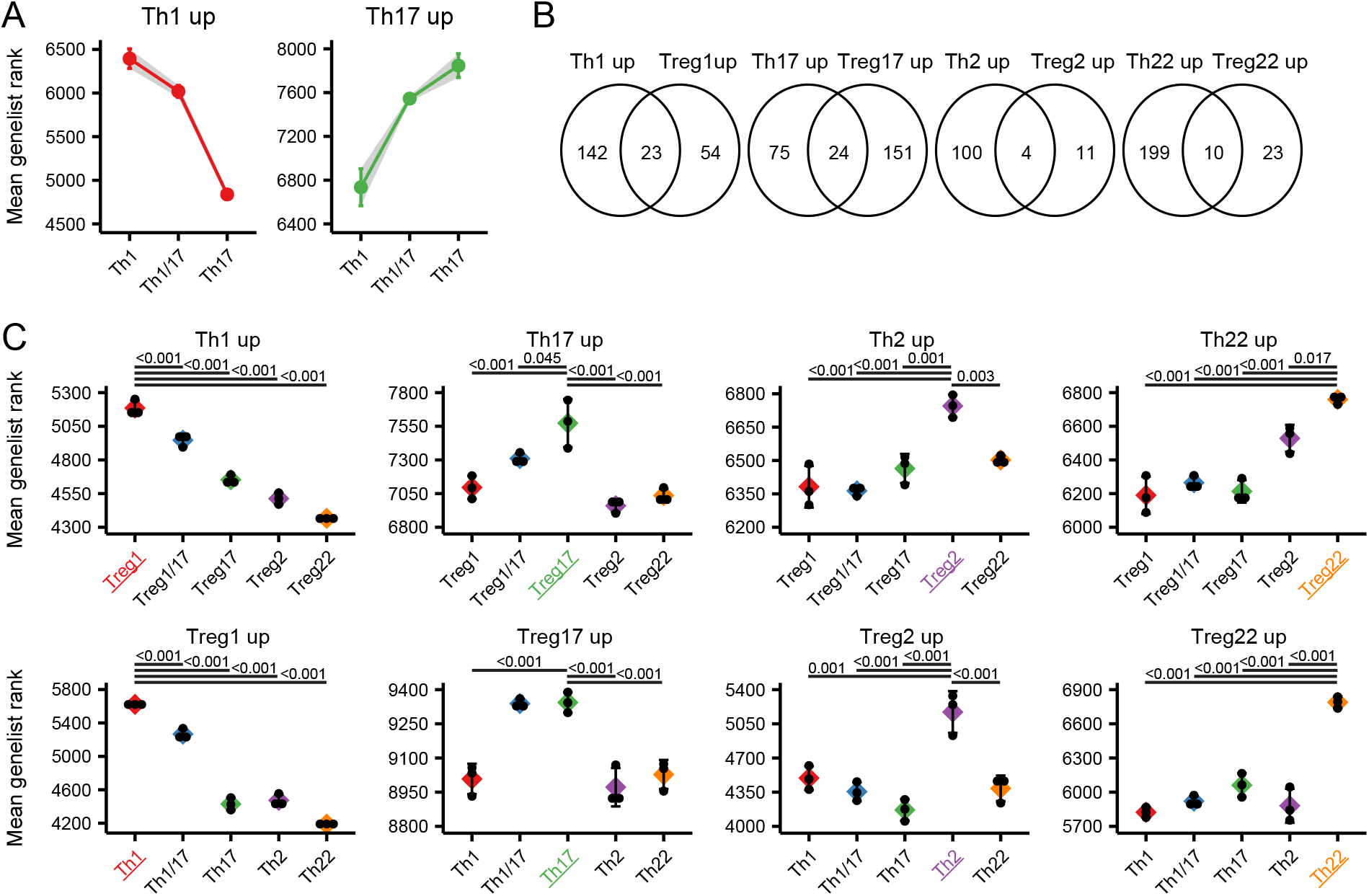
Shared transcriptional signatures of mirror Th and Treg cell populations. **A)** Mean genelist rank of the Th1/17 subset compared to the Th1 and the Th17 subset examining genes significantly upregulated in the Th1 (left) and Th17 (right) populations. **B)** Venn diagram showing the overlap in genes specifically upregulated in mirror Th and Treg cell populations. **C)** Mean genelist rank of Th signature genes in each Treg cell population (top), or mean genelist rank of Treg signature genes in each Th cell population (bottom) as indicated. Significance determined by ANOVA followed by Tukey HSD, and significantly different comparisons with the population of interest (highlighted with color) are indicated.

### Enrichment of Th and Treg transcriptional signatures in bulk RNA-seq data

The identification of specific transcriptional signatures for each of the Th and Treg cell subsets from peripheral blood raises the possibility that the genelist rank approach could be used to functionally characterize bulk Th and Treg samples. To test this idea, we first applied the Treg subset signatures to ranked genelists of RNA-seq samples from peripheral blood Treg cells that were sorted into 4 populations based on their expression of the cytokines IFN-γ and IL-10 (Sumida et al., 2018). As expected based on their preferential production of these cytokines, we found that genes specifically upregulated in Treg1 cells were significantly enriched in the IFN-γ^+^IL-10^+^ and IFN-γ^+^IL-10^−^ cells, that Treg17 signature genes were also enriched in the IFN-γ^−^ IL-10^+^ cells, and that Treg2 and Treg22 signature genes were elevated in the IFN-γ^−^IL-10^−^ cells (Fig 7A). To further extend this approach, we applied our signatures to data from bulk Th and Treg cells isolated from either peripheral blood or from human breast carcinoma (Plitas et al., 2016). In this case, Th cells from tumors were enriched in Th1 signature genes, whereas those from PBMC were comparatively enriched in Th17 and Th2 signature genes. This is consistent with the initial characterization of these data reporting high expression of Th1 signature genes *EOMES, GZMK, CXCR3* and *IFNG* in tumor-infiltrating Th cells (Plitas et al., 2016). Similarly, Treg cells from the tumors were enriched in Treg1 signature genes, but showed decreased expression of Treg17 and Treg22 signature genes. Lastly, we applied the Th cell signatures to RNA-seq data comparing gene expression by CCR7^+^ central memory T (T_CM_) cells and CCR7^−^ effector memory T (T_EM_) Th cells that express the skin homing receptor cutaneous lymphocyte antigen (CLA) from peripheral blood with CLA^+^ T_CM_ and T_EM_ from skin. In this analysis, we found a significant enrichment of Th22 signature genes in both T_CM_ and T_EM_ populations from skin. Th22 cells in the blood express a set of surface adhesion molecules and chemokine receptors indicative of skin-tropism, including CLA, CCR4 and CCR10, and IL-22-producing T cells are enriched in the skin where this cytokine can act directly on keratinocytes to induce tissue-repair and anti-microbial responses (REF). Together, these examples demonstrate the utility of the gene signatures we have identified for functional analysis of bulk Th and Treg cells isolated from both healthy and diseased tissues.

**Figure 7:**
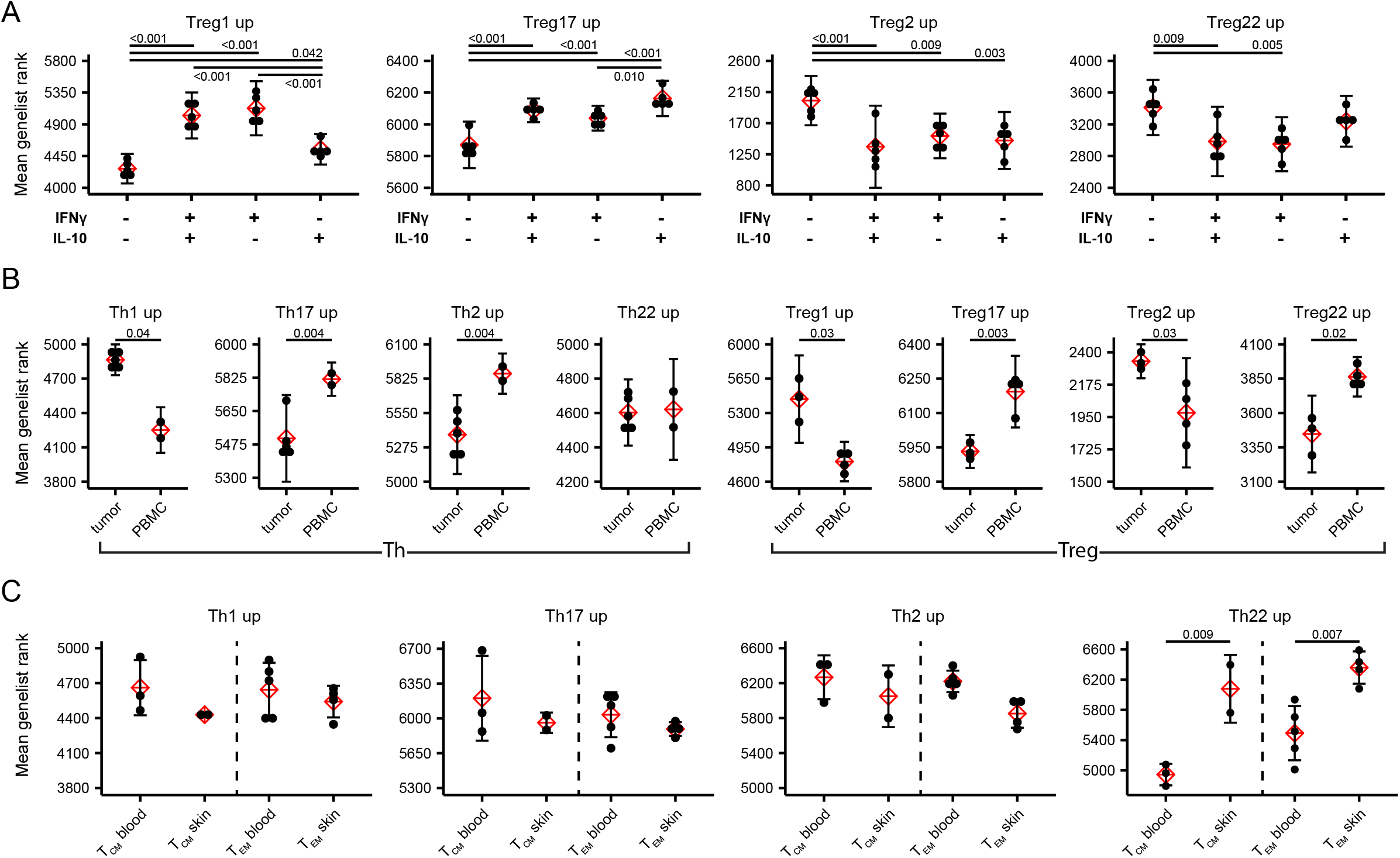
Predictive use of Th and Treg population signatures in bulk RNA-seq data. **A)** Mean genelist rank of each of the Treg population signatures was applied to RNA-seq data from Tregs sorted on the basis of IFN-γ and IL-10 production as indicated (data from GEO dataset GSE116283). Significance determined by ANOVA followed by Tukey HSD. **B)** Mean genelist rank of each of the Th and Treg population signatures was applied to RNA-seq data from Th and Treg cells sorted from tumor or peripheral blood of patients with breast carcinoma (data from GSE89225). Significance determined by Student t-test. **C)** Mean genelist rank of each of the Th population signatures was applied to RNA-seq data from CLA^+^CCR7^+^ T_CM_ and CLA^+^CCR7^−^ T_EM_ sorted from blood and skin (data from GSE149090). Significance determined by Student t-test.

## Discussion

The functional specialization of CD4^+^ T cells is the cornerstone of effective adaptive immune responses that prevent the growth and spread of various pathogens while limiting collateral tissue damage and autoimmunity (Mahnke et al., 2013). Through analysis of 11 human CD4^+^ T cell populations sorted directly *ex vivo*, we have comprehensively defined the transcriptional basis for human CD4^+^ T cell specialization, resulting in several important and novel insights. We assessed the transcriptional relationship between the various CD4^+^ T cell populations, and defined core Th and Treg gene signatures as well as unique signatures of each population that can be applied to analysis of existing RNA-seq data sets for functional profiling of human Th and Treg cells in different tissue sites.

The majority of Treg cells are believed to develop in the thymus upon encounter with self-antigen, and therefore this can be thought of as the first branch point in the functional differentiation of CD4^+^ T cells. At barrier sites, Treg cells can also develop from conventional naïve T cells activated in tolerogenic conditions to help enforce tolerance to harmless environmental antigens, including components of the skin and intestinal microbiomes (Lee et al., 2011). The fact that all of the Treg cell populations differentially expressed genes generally involved in T cell activation and proliferation is indeed consistent with each containing large fractions of highly reactive cells. Among these, it is interesting that each Treg population showed increased expression of the transcription factor TOX, which has recently been found to promote and exhausted phenotype in chronically stimulated CD8^+^ T cells and thus to prevent activation induced cell death (Alfei et al., 2019; Khan et al., 2019; Scott et al., 2019).

Interestingly, despite displaying similar phenotypic diversity, transcriptionally the Treg populations were far more uniform then their Th counterparts. This likely reflects the dominant role of Foxp3 on the Treg transcriptome, and the ability of this single factor to convert potentially pathogenic effector T cells into potent Treg cells (Hori et al., 2003). Induction of Foxp3 in Treg cells initiates dramatic changes in gene expression that underlie their suppressive functions. Foxp3 activates a broad transcriptional program that controls Treg cell function and homeostasis, and includes genes such as *CTLA4* and *IL2RA* (CD25). Foxp3 also inhibits expression of key effector cell molecules such as *IL2, IFNG,* and *CD40LG* that were more highly expressed in Th cells (Bhairavabhotla et al., 2016). Indeed, our analysis clearly shows that despite these shared properties, functionally specialized Th subsets are transcriptionally distinct from their Treg cell mirror counterparts, and we used each of these comparisons to identify ‘core’ Th and Treg gene signatures that confirm and extend previous analysis of the specific Treg cell transcriptome. All Treg populations showed increased expression of core immunosuppressive genes (*CTLA4, TIGIT, ITGB8, LRRC32, ENTPD1)* and specific chemokine/cytokine receptors (*CCR3*, *CXCR6*, *CSF2RB, IL1R1*). Unlike Th cells that undergo extensive functional diversification based on differential expression of key effector cytokines, most of the key functional immunoregulatory molecules expressed by Treg cells were expressed to some degree by all of the Treg cell populations. Thus, the phenotypic diversity observed among Treg cell populations may function primarily to direct Treg cells with the appropriate specificities to sites of Th1, Th2, Th17 or Th22 mediated inflammatory responses (Campbell and Koch, 2011).

The phenotypic and functional diversity of Th cells has been extensively studied. Although initially thought of as terminally differentiated subsets, it has become increasingly clear that there is significant plasticity among different Th populations (Bonelli et al., 2014). In particular, there is substantial plasticity in Th17 cells, which can adopt a hybrid Th1/17 phenotype in response to inflammatory cytokines such as IL-12 or IL-1β (Duhen and Campbell, 2014; Sallusto et al., 2012). This trans-differentiation is most relevant in the context of autoimmune and inflammatory diseases such as multiple sclerosis and Crohns disease in which hybrid populations and/or ‘ex-Th17’ cells have been implicated in pathogenesis (Hirota et al., 2011; Ramesh et al., 2014). Indeed, our transcriptional analysis of Th1/17 cells showed that they are hybrid population that shares the transcriptional signature of both Th1 and Th17 cells. This includes not only shared expression of *RORC* and *TBX21*, but also the highest levels of the cytokine receptors *IL23R* and *IL12RB2* and some unique genes such as the multi-drug resistance transporter *ABCB1* and the inhibitory receptor *KLRB1* (CD161). Similarly, Treg1/17 cells transcriptionally resembled both the Treg1 and Treg17 populations, indicating that there may be significant phenotypic plasticity among these populations as well.

The identification and isolation of distinct Th and Treg populations based on differential chemokine receptor expression works well in CD4^+^ T cells isolated from healthy human blood. However, similar analyses from non-lymphoid tissues can be challenging due to changes in phenotype that occur upon cellular entry into certain tissue sites, cleavage of specific markers during enzymatic digestion required for isolation of T cells from some non-lymphoid tissues, and the small number of T cells obtained from clinical samples and tissue biopsies. Our identification of unique subset-specific transcriptional signatures provides the opportunity to re-analyze bulk RNA-seq data of tissue Th and Treg cells to determine if any of these signatures are enriched. We validate the utility of this approach in 3 independent data sets derived from cytokine producing Treg cells from peripheral blood, Th and Treg cells infiltrating breast carcinomas, and Th cells found in the skin. Thus, these signatures can be used to guide analyses of both bulk and single-cell RNA-seq data to assess Th and Treg cell specialization. This will be particularly useful in analyses of small clinical samples that are limiting for conventional flow cytometric or functional analyses.

Our comprehensive analysis of human CD4^+^ T cells provides a new framework for understanding the transcriptional basis of Treg and Th specialization, and an important resource for analysis of transcriptomic data from T cells in healthy and diseased tissues. Cross-referencing these transcriptional profiles with analyses of epigenetic modification and transcription factor binding in different Treg and Th populations will help to further define the molecular mechanisms that underlie the diversity of CD4^+^ T cells, and suggest new ways to manipulate specific pathways to tune Th or Treg responses in cancer, autoimmunity and chronic infection.

